# SARS-CoV-2 infection studies in lung organoids identify TSPAN8 as novel mediator

**DOI:** 10.1101/2021.06.01.446640

**Authors:** Lisiena Hysenaj, Samantha Little, Kayla Kulhanek, Oghenekevwe M. Gbenedio, Lauren Rodriguez, Alan Shen, Jean-Christophe Lone, Leonard C. Lupin-Jimenez, Luke R. Bonser, Nina K. Serwas, Kriti Bahl, Eran Mick, Jack Z. Li, Vivianne W. Ding, Shotaro Matsumoto, Mazharul Maishan, Camille Simoneau, Gabriela Fragiadakis, David M. Jablons, Charles R. Langelier, Michael Matthay, Melanie Ott, Matthew Krummel, Alexis J. Combes, Anita Sil, David J. Erle, Johannes R. Kratz, Jeroen P. Roose

## Abstract

SARS coronavirus-2 (SARS-CoV-2) is causing a global pandemic with large variation in COVID-19 disease spectrum. SARS-CoV-2 infection requires host receptor ACE2 on lung epithelium, but epithelial underpinnings of variation are largely unknown. We capitalized on comprehensive organoid assays to report remarkable variation in SARS-CoV-2 infection rates of lung organoids from different subjects. Tropism is highest for TUBA- and MUC5AC-positive organoid cells, but levels of TUBA-, MUC5A-, or ACE2-positive cells do not predict infection rate. We identify surface molecule Tetraspanin 8 (TSPAN8) as novel mediator of SARS-CoV-2 infection, which is not downregulated by this specific virus. TSPAN8 levels, prior to infection, strongly correlate with infection rate and TSPAN8-blocking antibodies diminish SARS-CoV-2 infection. We propose TSPAN8 as novel functional biomarker and potential therapeutic target for COVID-19.

## Introduction

Severe Acute Respiratory Syndrome Coronavirus-2 (SARS-CoV-2) has caused a global pandemic of coronavirus disease (COVID-19) with over 170M cases worldwide (https://coronavirus.jhu.edu/map.html) as of May 2021. Clinically, SARS-CoV-2 can lead to lethal acute respiratory distress syndrome (ARDS) ^1^. Many SARS-CoV-2 studies have focus on the immune system, currently, we do not understand how the lung epithelium itself may play a role in variation in disease spectrum of COVID-19.

The lung airway epithelium defends against pollutants, allergens, and pathogens and is composed of a variety of cell types including basal cells, ciliated cells, mucus-producing goblet cells, secretory cells, and neuroendocrine cells. The distal lung epithelium also includes alveolar type 1 (AT1) cells, which mediate gas exchange, and AT2 cells, which secrete surfactant ^2, 3^. In the case of COVID-19, studies thus far suggest that SARS-CoV-2 infects mostly ciliated cells, goblet cells, AT2 cells but also basal stem cells ^4, 5, 6, 7, 8^. Comprehensive SARS-CoV-2 infection studies focusing on lung epithelial cell subsets in different individuals have not been reported to date.

Coronavirus (CoV) particles are spherical with three viral proteins anchored in the envelope: the triple-spanning membrane (M) protein, the envelope (E) protein, and the spike (S) protein, which form the characteristic trimeric spikes ^9, 10, 11^. The surface spike glycoprotein of SARS-CoV-2 binds to human ACE2 (Angiotensin-converting enzyme 2) ^12^. Binding of the S protein to ACE2 mediates membrane fusion and viral entry. The S protein is cleaved by host cell type II trans-membrane serine proteases resulting in spike protein activation and viral entry ^13, 14, 15, 16^. As such, ACE2 and Transmembrane protease, serine 2 (TMPRSS2) are critical for SARS-CoV-2 entry into the cell ^17^.

Viruses typically co-opt various host proteins to maximize infectious potential ^18^. The wide variation in SARS-CoV-2 infection rates and COVID-19 severity suggests that there must be facilitators other than ACE2 and TMPRSS-2 that have yet to be discovered. Epithelial organoids have gained traction as a physiological platform for personal medicine because these organoids retain patient-specific traits^19^. A handful of studies have used alveolospheres and organoids to study SARS-CoV-2 epithelial response ^20, 21, 22, 23, 24, 25^ and tropism ^7, 26, 27, 28, 29, 30^. A comprehensive analysis of SARS-CoV-2 infection in a large panel of lung organoids has not been reported to date. Here, we performed a comprehensive analysis with the goal of obtaining insights for different subjects as well as discovery of new molecules that play a role in infection.

## Results

### A comprehensive lung organoid biobank

We generated organoids from biopsies from healthy donors or adjacent normal tissue from lung cancer patients undergoing surgery (**Table 1**). 3D lung organoids from twenty subjects were expanded through passaging and cryopreserved, generating a biobank (**Figs. 1A and 1B, and Suppl. Fig S1A and S1B**).

**Figure 1:**
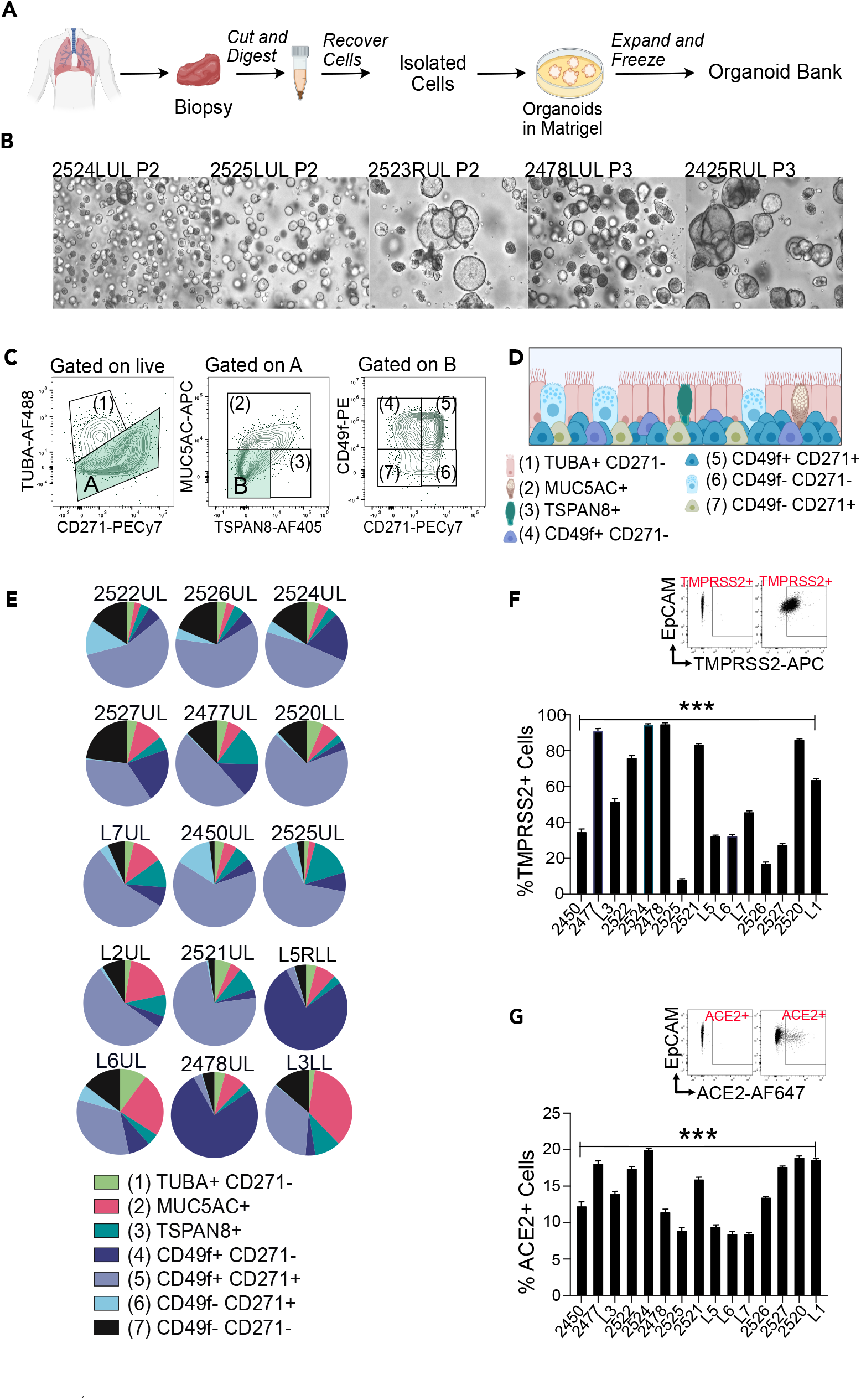
Donor-derived lung organoids are stable and distinctive. A) Workflow of lung organoid generation. B) Brightfield images of lung organoids derived from different donors. C) Spectral Flow Cytometry gating strategy to define seven cell populations: (1)TUBA+ CD271- as ciliated cells. (2) MUC5AC+ TUBA- as goblet-like cells. (3) TSPAN8+ MUC5AC-TUBA- as pre-goblet cells. (4) CD49f+ CD271+, (5) CD49f-CD271+, (6) CD49f+ CD271+ as basal cells and (7) CD49f-CD271- as undefined cells. D) Scheme of cell types found in the 3D lung organoids. E) Pie charts representing spectral flow cytometry analyses of lung organoids from indicated donors. F-G) Percentage of F) ACE2+ and G) TMPRSS2+ cells in lung organoids based on spectral flow cytometry analyses. Bars represent mean, error bars are SEM, n=3. Paired t-test, *p<0.05; ns, non-significant.

Spectral flow cytometry analyses with single tube staining of fourteen markers (termed spectral flow here), including a panel of recently developed airway epithelial markers ^31^ revealed diversified compositions of cell subsets in fifteen lung organoids. We identified ciliated *like* cells (TUBA^high^ CD271^neg^) ^32^, goblet *like* cells (TUBA^neg^, MUC5AC^+^)^33^, pre-goblet *like* cells (TUBA^neg^, MUC5AC^−^,TSPAN8^+^)^34^, three populations of cells expressing basal cell markers CD49f^+^CD271^+^, CD49f^neg^CD271^+^, CD49f^+^CD271^+^, and a population of CD49f^neg^CD271^neg^TUBA^neg^MUC5AC^neg^TSPAN8^neg^ cells (**Figs.1C and 1D**). Lung organoids derived from different donors displayed distinctive cell type compositions (**Fig. 1E**). By contrast, compositions of organoids generated from upper lobe and lower lobe samples from the same patient were very similar (**Suppl. Fig S1B**) and analysis of different passages from distinct organoids demonstrated that our lung organoids are stable and retain patient-specific composition in passaging (**Suppl. Fig S1C**). Spectral Flow Cytometry staining for intra-cellular TMPRSS2 (**Fig. 1F**) and extracellular ACE-2 (**Fig. 1G**) allowed for the assessment of the percent of cells expressing these proteins that play critical roles in SARS-CoV-2 entry. Pearson correlations between age and the proportions of goblet *like-*, ciliated *like*-, and basal-cells in unmanipulated organoids were not significant (**Suppl. Fig S1D-S1F**). We selected a panel of twelve lung organoids that captured the variety in cell composition, age, sex, and ACE-2 and TMPRSS2 expression for further studies.

### Viral infections of lung organoids

We utilized H1N1/PR8 influenza virus to optimize viral infection of lung organoids (**Suppl. Fig. S3A-D**). Following infection, we assessed cell-composition changes with spectral flow, adding the cKit marker (**Suppl. Fig S2 and S3F**), as this receptor has been suggested to mark lung regeneration upon injury ^35, 36^ and tropism of cell types for H1N1 (**Suppl. Fig S3G**). Since viral infections trigger epithelial interferon responses, which help orchestrate immune responses ^37^, we also stained for co-stimulatory molecules CD80 and CD86 ^2, 38^, or immune-activating molecules CEACAM5 and CEACAM6 ^39, 40^. ACE2 has recently been described as an interferon-upregulated gene ^41^. We observed upregulation of these five cell surface molecules after H1N1/PR8 infection (**Suppl. Fig. S3H**).

Having established consistent infection with H1N1/PR8, we switched to SARS-CoV-2 (**Fig 2A**). Whole mount organoids revealed the presence of double stranded RNA (dsRNA) and viral nucleocapsid protein (N) in the infected lung organoids, that were also stained for CD49f and acTUBA (**Fig. 2B**). Five independent SARS-CoV-2 infections of lung organoid 2522UL demonstrated that our infections were robust and consistent (**Suppl. Fig S4A-D**). Spectral flow of dsRNA staining, marking replicating virus, revealed a remarkable variation in the percentage of (replicating) SARS-CoV-2-positive cells in twelve different organoids and 31 SARS-CoV-2 infections with means ranging from fifteen to zero percent (**Figs. 2C and 2D**). The percentage of cell death in the organoids (**Suppl. Fig S5A**) and infection rates (**Fig. 2D**) showed different patterns. We characterized cell compositions (**Fig. 2E** and **Suppl. Fig S5B**) and plotted changes using PCA (principal component analysis) (**Suppl. Fig S5C**). SARS-CoV-2 infection resulted in increased proportions of cells expressing acTUBA (**Fig. 2F**) and cKit (**Fig. 2G**), but no alterations in the fraction of basal cells (CD49+ or CD271+), MUC5AC-positive, or CD49f^neg^CD271^neg^ cells (**Fig. 2H,Suppl. Fig S5B** and **S5D**).

**Figure 2:**
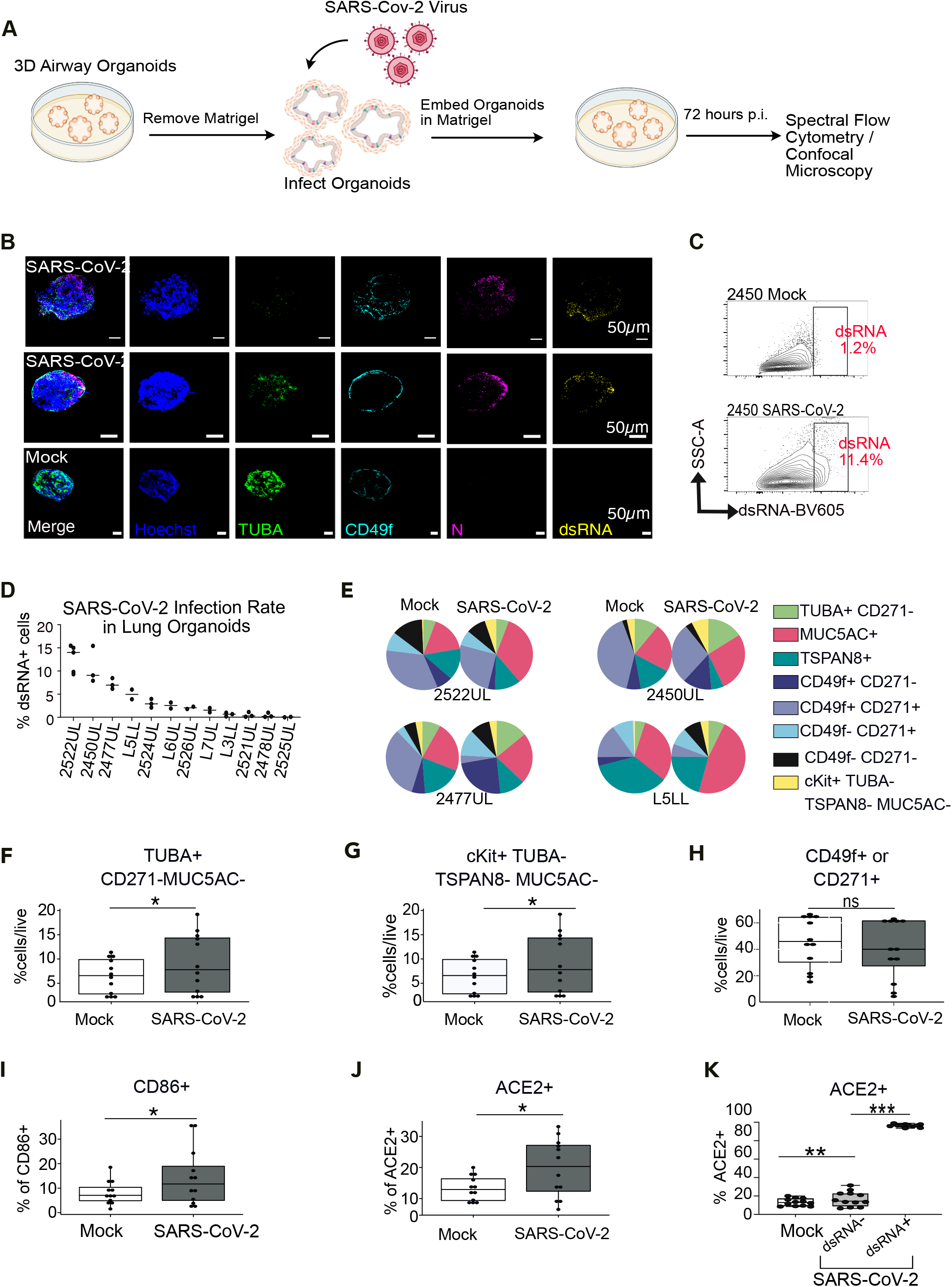
Patient derived organoids show distinct responses to SARS-CoV-2 infection. A) Experimental scheme SARS-CoV-2 (MOI=0.3) infection. B) Representative, confocal images (Z-stack) of SARS-CoV-2 infected (top and middle panel) and mock infected (bottom panel) whole mounted organoids. Scale bars are 50uM. C) Spectral flow for dsRNA, marking cells with replicating SARS-CoV-2. D) Percentage of dsRNA-positive cells in SARS-CoV-2 infected lung organoids E) Composition of Mock and SARS-CoV-2 infected organoids. F-H) Comprehensive analysis of % of cell populations (F) Ciliated like cells (TUBA+), G) cKit+, H) Basal cells) in Mock and SARS-CoV-2 infected organoids for 12 different donor derived lung organoids. Each point represents the mean of % cell type for the distinct donor for Mock or SARS-CoV-2 conditions. Wilcoxon signed-rank paired test, *p<0.05, N=12, n=3. I-J) Percentages of ACE2+ and CD86+ cells. Each point represents the mean of % cell type for the distinct donor for Mock or SARS-CoV-2 conditions. Wilcoxon signed-rank paired test, *p<0.05, N=12, n=3. K) The percentage of ACE2+ cells in live cells of non infected organoids (Mock), and the percentage of ACE2 positive cells in live dsRNA- and in dsRNA+ live cells of SARS-CoV-2 infected organoids at 72h p.i.. Each point represents the mean of % cell type for the distinct donor for Mock or SARS-CoV-2 conditions. Friedman Test, **p< 0.01; ***p<0.001. ns, non-significant.

In terms of functional markers, CD86-positive cells were increased (**Fig. 2I**), but SARS-CoV-2 infection did not cause increases in proportions for CD80-, CEACAM5-, and CEACAM6-expressing cells (**Suppl. Fig S5E**), indicating that there was not a uniform induction of an interferon response program in all cells of infected organoids. Lungs from COVID-19 patients demonstrate elevated ACE-2 protein expression ^42^. We observed significant increases in the proportions of ACE2-positive cells (**Fig. 2J**) upon SARS-CoV-2 infection in the 12 lung organoids. The spectral flow allows us to compare mock-infected to SARS-CoV-2 exposed/infected (dsRNA+), and to SARS-CoV-2 exposed/uninfected (dsRNA-). These comparisons revealed that nearly one hundred per cent of dsRNA-positive, infected cells were ACE2-positive (**Fig. 2K**). Similarly, we observed high proportions of cells positive for CD80, CD86, CEACAM5, and CEACAM6 when analyzing exposed/infected specifically (**Suppl. Fig S4E**).

### Finding the rules of variation in SARS-CoV-2 infection

The lung organoids from subject 2525 express low TMPRSS2 (**Fig. 1G**), providing an explanation for the low infection rate in 2525 (**Fig. 2D**). For the other organoids an explanation for the variation was lacking. Our panel of twelve organoids and 31 SARS-CoV-2 infections allowed us to explore if there are rules that help explain the infection variation that ranged from thirteen to zero percent on average. We observed a tropism of SARS-CoV-2 for acTUBA- and MUC5AC- positive cells (**Fig. 3A**). Depicting PCA of many factors in a circle of correlation, we established that there are probable correlations between amount ciliated cells (acTUBA positivity), ACE2 positivity, and CD86 positivity, determined in non-infected organoids and subsequent infection rate. There is also a probable correlation between the age of the donor and infection, since the arrows are of substantial length and point in the same direction (**Fig. 3B**). However, statistical analysis of age (**Fig. 3C,** Pearson 0.45, Pv = 0.14), TUBA-positivity (**Fig. 3D**, Pearson 0.48, Pv = 0.11), and CD86-, cKit-, or MUC5AC- positivity (**Suppl. Fig S5F**) as single factors revealed that none correlate with SARS-CoV-2 infection rate in organoids.

**Figure 3:**
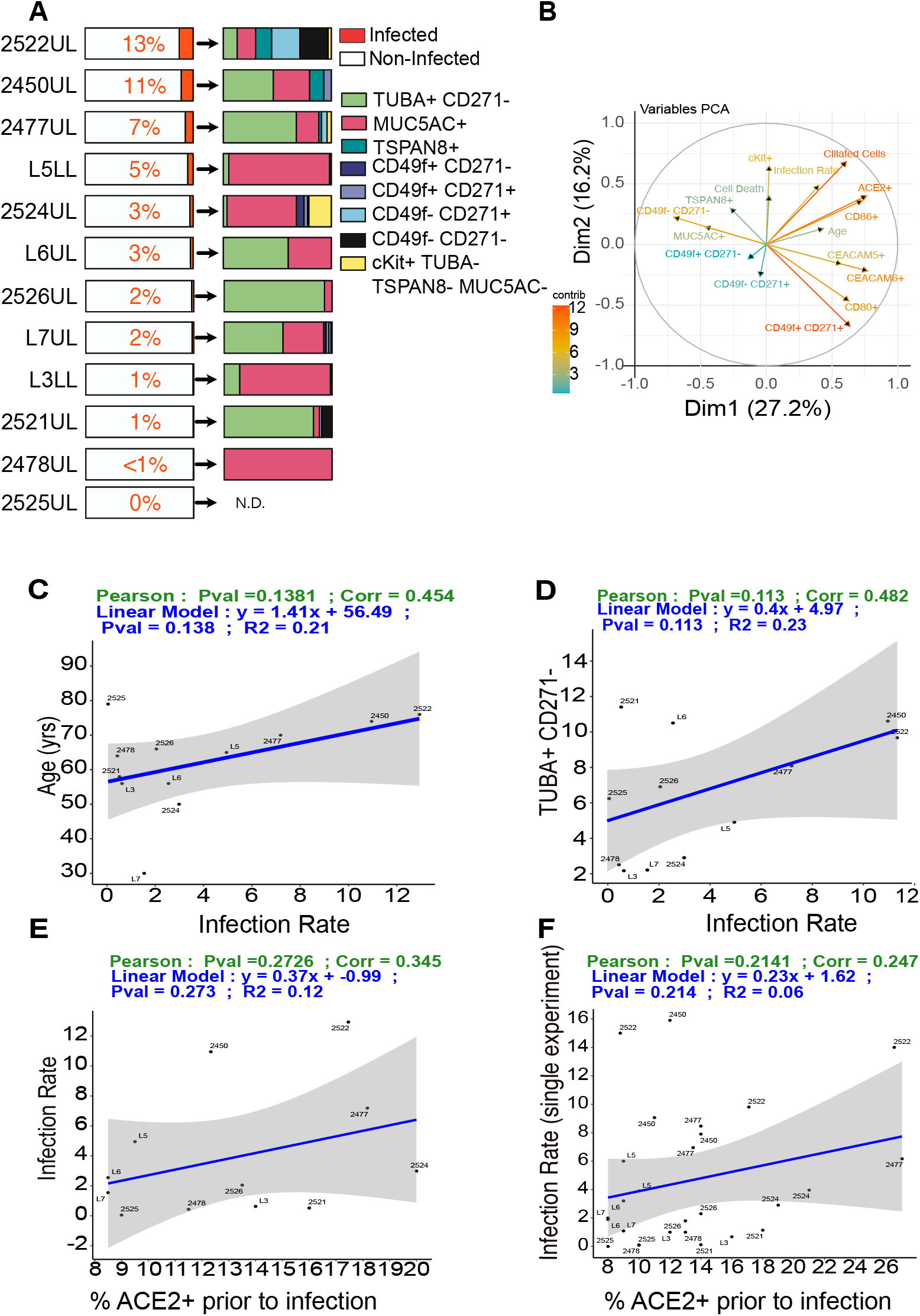
Susceptibility of lung organoids to SARS-CoV-2 infection is not predicted by ACE2. A) Percentage of dsRNA+ cells (left) and the cell types in dsRNA+ cells (right) in SARS-CoV-2 infected organoids (MOI=0.3, 72h p.i.). Mean values of 31 infections are depicted and plotted. B) PCA of different variables impacting infection. Each arrow corresponds to one biological descriptor; the longer the arrow, the better the representation (the color displays the cos2). C) Linear regression modeling the relationship between age of the donor and infection rate. D) Linear regression modeling the relationship between percentage of TUBA+ CD271+ cells and infection rate. Each point represents the mean of % cell type for the distinct donor. E) Linear regression modeling the relationship between Infection rate and % ACE2+ cells prior to infection. Each point represents the mean of % cell type for the distinct donor. F) Infection rate and % ACE2+ cells prior to infection for 31 single infections. Statistical significance is displayed by P values of Pearson Correlation for each graph. Mean values, N=12, n=3.

The percentage of ACE2 positive cells was significantly upregulated upon infection of lung organoids (**Fig. 2J**) and nearly all dsRNA-positive cells are also ACE2-positive (**Fig. 2K**). However, the proportion of ACE2-positive cells in the organoids prior to infection, either as mean infection or as single datapoints of all 31 infections, did not correlate with infection rate (**Fig. 3E and 3F**). These findings motivated us to search for novel molecules that show strong correlations with rate of infection and may function as host proteins co-opted by SARS-CoV-2 in infecting the lung.

### Single cell gene expression points to TSPAN8

In search of new mediators of SARS-CoV-2 infection, we infected four organoids with SARS-CoV-2 and performed single-cell RNA sequencing (sc-RNAseq) of organoids. Unsupervised clustering analysis based on most variable gene expression across all cells ^43^, regardless of infection status, identified sixteen unique cell subsets represented in a UMAP plot (**Fig. 4A**). Based on the top five most differentially expressed genes by cluster we assigned relative identities to these sixteen populations (**Suppl. Fig S6**), though it should be noted that assignment of identity of single cells of different lung organoids was not our objective here. Furthermore, while clustering analysis of individual organoids showed significant differences (**Suppl. Fig S7A**), read counts were relatively low for organoids L7 and 2524. For this reason, we treated the four organoids as one collective dataset, while capitalizing on the resolution of the sc-RNAseq for discovery.

**Figure 4:**
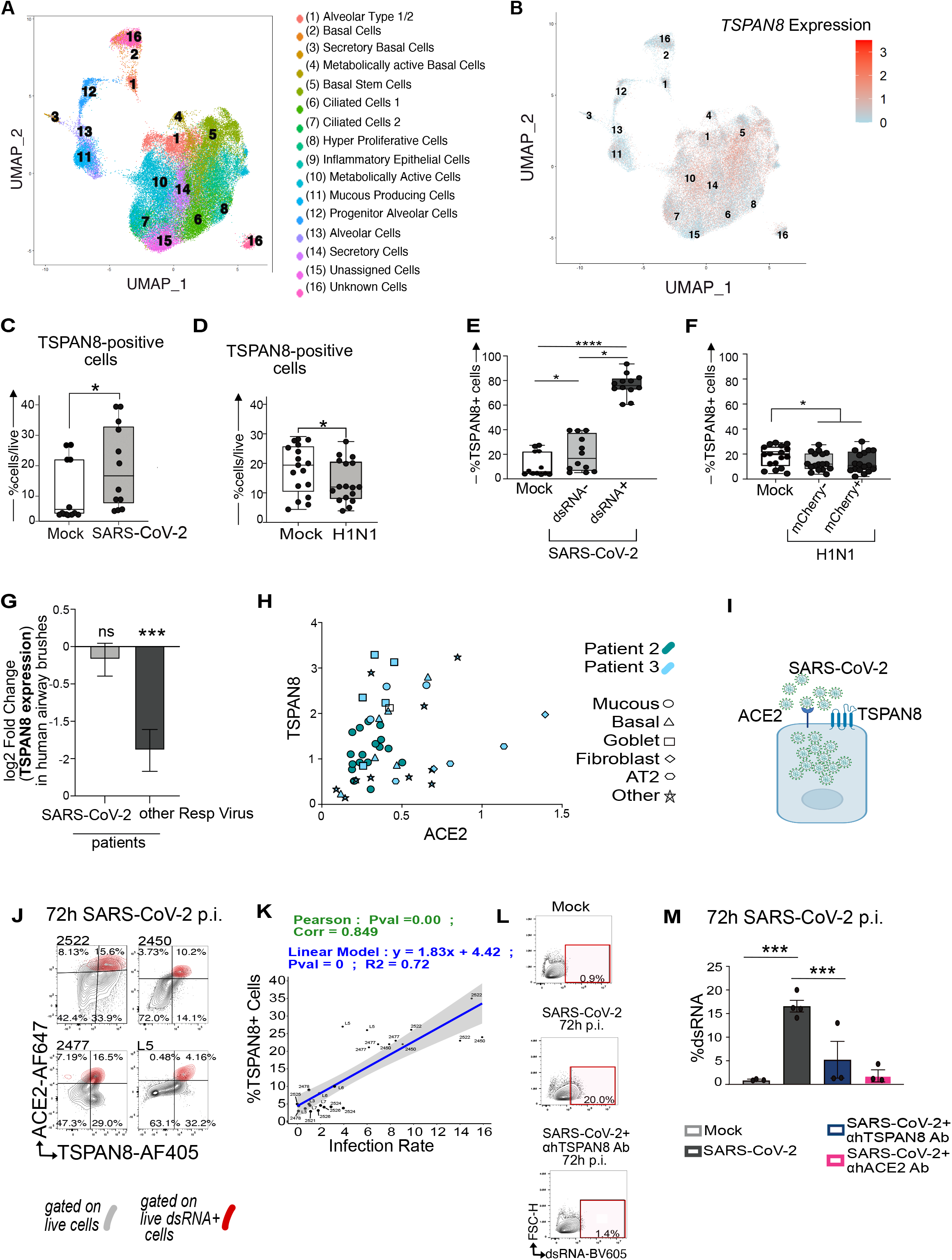
TSPAN8 is a facilitator, risk factor, and target for SARS-CoV-2. A) UMAP reduction on the merged cell data with overlaid clusters and identified cell types. B) TSPAN8 expression overlay on the UMAP seen in (A). C-D) Percentage of TSPAN8 positive cells for 12 SARS-CoV-2 (MOI=0.3) infected or 3 H1N1(MOI=0.15) infected lung organoids for Mock and infected organoids at 72h p.i.. Every point represents a single experiment. Each experiment was performed indipensently at least 2 times using three replicates per experiment. Friedman Test, *p<0.05; ****p<0.0001. E-F) The percentage of TSPAN8+ cells in live cells of non infected organoids (Mock), and the percentage of ACE2 positive cells in (E,F) live dsRNA- and in dsRNA+ live cells of SARS-CoV-2 infected organoids or mCherry+ and mCherrry-live cells of H1N1 infected organoids at 72h p.i.. Each point represents the mean of % cell type for the distinct donor for Mock or viral infected conditions. Friedman Test, **p<0.01; ***p<0.001. ns, non-significant. H) Analyses of ACE2 and TSPAN8 co-expressing cells in vivo in normal human lung, from 10X single-cell sequencing data from Travaglini et al. I) SARS-CoV-2 virus, ACE2, and TSPAN8 in a representation of lung epithelial cell G) Differential expression of TSPAN8 in nasal swabs of adult patients with acute respiratory illness (ARI) due to COVID-19 (n=93) or other viral infection (n=41), in comparison to patients with ARI due to non-viral etiology (n=100) J) Spectral flow cytometry plots showing live dsRNA-cells (in gray) and live dsRNA+ cells (overlaid in red) in four different SARS-CoV-2 infected organoids at 72h p.i. K) Correlation between percentage of TSPAN8 positive cells prior to infection and SARS-CoV-2 infection rate. 31 individual infections. P values indicate statistical significance. L) Spectral Flow Cytometry Plots of dsRNA-positive cells in Mock, SARS-CoV-2 infected organoids in presence or absence of TSPAN8 blocking antibody. M) Percentage of of dsRNA+ cells in Mock, SARS-CoV-2 infected organoids in presence or absence of TSPAN8 or ACE2 blocking antibody. SARS-CoV-2 infection in the indicated blocking conditions (N=1, n=3). M) Percentage of dsRNA-positive cells in Mock, SARS-CoV-2 infected organoids in presence or absence of TSPAN8 or ACE2 blocking antibody. SARS-CoV-2 infection in the indicated blocking conditions (N=1, n=3). Bars represent mean, error bars are SEM, n=3. Paired t-test, *p<0.05; ns, non-significant.

We next focused on cells expressing SARS-CoV-2 transcript to identify specific transcriptomic signatures in infected cells (**Table 2**). Metascape pathway analysis ^44^ confirmed a signature of inflammatory response, in agreement with the identity of virus-infected epithelial cells (**Suppl. Fig S7C**). We plotted a cluster of sixty genes that were differentially expressed in single cells positive for SARS-CoV-2 viral read compared to false identities (**Suppl. Fig. S7B** and **Table 3A**). We mined our dataset in search of possible novel receptors for SARS-CoV-2 and noted that the surface molecules TSPAN8, AREG, and CD24 (**Suppl. Fig. S7B** and **Table 3B**) were positively expressed in single cells containing SARS-CoV-2 reads. We were particularly intrigued by TSPAN8 (Tetraspanin 8), since members of the TSPAN family are believed to promote cell entry of different viruses ^45^, but TSPANs have not been linked to SARS-CoV-2. Single cell *TSPAN8* mRNA reads were present in 64% of cells with SARS-CoV-2 reads (**Suppl. Fig. S7B**) and *TSPAN8* expression was not limited to a unique cell subset (**Fig. 4B**).

### TSPAN8 facilitates SARS-CoV-2 infection and can be inhibited

*TSPAN8* mRNA expression had been documented in the supplemental dataset of a single SARS-CoV-2-infected lung organoid ^8^, but the role of TSPAN8 in SARS-CoV-2 infection has not been investigated. Fortuitously, we had included anti-TSPAN8 antibody in our spectral flow to distinguish goblet like cells from pre-goblet cells (**Suppl. Fig. S2)**, allowing for surface protein expression to confirm the sc-RNAseq discovery. The proportion TSPAN8-positive cells in the organoids increased upon SARS-CoV-2 infection (**Fig. 4C)**, but decreased with H1N1 infections (**Fig. 4D**). Just as for ACE2 (**Fig 2K**), most infected cells (dsRNA positive) expressed TSPAN8 (**Fig. 4E**), a pattern not observed for H1N1 (**Fig. 4F**). We further investigated *TSPAN8* in the context of viral infections. Nasal swabs of adult patients with acute respiratory illness (ARI) ^46^ due to non-COVID-19, respiratory viral infection revealed downregulation of *TSPAN8* (**Fig. 4G**). However, such airway brushes from COVID patients revealed that *TSPAN8* expression remained of the same level compared to comparison to patients with ARI due to non-viral etiology **Fig. 4G**.

Mining published scRNAseq data of human lungs ^3^ that predominantly reported on immune cell composition, we noted that of 61,000 cells sequenced from lung tissue (as opposed to blood), only 48 cells had expression of both TSPAN8 and ACE2 and all of these 48 cells were of epithelial origin (**Fig. 4H**). Furthermore, back-gating on dsRNA-positive cells and overlaying this onto a ACE2/TSPAN8 contour plot, our spectral flow revealed that a large proportion of SARS-CoV-2 infected cells simultaneously expresses both ACE2 and TSPAN8 (**Fig. 4J** and **Suppl. Fig S7F**). Collectively, these data suggest that the two surface proteins may cooperate in SARS-CoV-2 infection (**Fig. 4I**).

Given the TSPAN8 data and the report that TSPAN family member CD9 facilitates MERS-CoV infection in mice ^47^, we considered whether TSPAN8 is a facilitator of SARS-CoV-2 infection (**Fig. 4I**). First, we explored TSPAN8 as biomarker. Different from our studies on ACE2 expression (**Figs. 3E** and **3F**), high percentages of TSPAN8-positive cells prior to infection did correlate very strongly with subsequent SARS-CoV-2 infection rates in the 31 infections (**Fig. 4K, Suppl. Fig S7G**). Lastly, we explored TSPAN8 as potential therapeutic target. We chose organoid 2450UL that displays high baseline TSPAN8 expression (**Suppl. Fig. S7F),** infected with SARS-CoV-2, and subjected these to blocking antibody treatment. Addition of a blocking antibody that binds to one of TSPAN8’s extracellular loops reduced SARS-CoV-2 infection by 60 per cent, roughly in the same range of inhibition of an ACE2-blocking antibody (**Figs. 4L and 4M**, **Supplemental Fig. S7H and I**), which had been reported previously ^14^.

## Discussion

Here we characterized the composition of 15 lung organoids derived from different donors. Multi-parameter spectral flow of cell sub-sets shows that the organoids are stable, but unique from subject to subject, generating a diverse lung organoid biobank. Subject-specific traits can be investigated, such as the pattern of decreasing amounts of mucus-producing cells as the age of the donor subject increases (**Suppl. Fig S1F**). H1N1/PR8 influenza and SARS-CoV-2 infection and spectral flow allowed for discovery of cell responses. Fifteen antibody spectral flow generates gating strategies where cell populations are embedded in each other, providing quantitative, linked data that can be analyzed in many ways. We were intrigued by the increase in cKit-positive cells after infection, as cKit has been suggested as marker of lung regeneration upon injury ^35, 36^. Possibly infected organoids sense damage and attempt to repair injury. Spectral flow-assisted gating on SARS-CoV-2-infected cells revealed a clear tropism for ciliated and MUC5AC-positive cells, but other cell types could also get infected. It should be noted here that our lung organoids are all embedded in Matrigel as we have not yet been able to obtain high throughput for organoid with air-liquid interface ^48^.

Neutralizing antibodies from COVID-19 patients have multiple targets ^49, 50^, suggesting that protective immune responses occur that may block interactions of molecules other than the S protein-ACE2 receptor pair. In our scRNA data *RALA* and *CD24* reads were present in infected cells. RALA (RAS like Proto-Oncogene A) was identified in the host-coronavirus protein network ^51^ and infection levels of a hepatoma cell line is reduced when deleted by CRISPR^17^. CD24 is a glycosyl-phosphatidyl-inositol (GPI)-anchored membrane protein that can repress the host response to DAMPS ^52^ and EXO-CD24 exosomes are explored in clinical trials as therapy for COVID19 patients (Clinical Trial NCT047477574). Studies using cell lines had also implicated additional mediators of infection, including AXL ^53^ CD147 ^54, 55^ and neuropilin-1 (NRP1) ^56, 57^.

We discovered that TSPAN8 strongly correlates with infected cells and infection rate. Tetraspanin proteins have four transmembrane domains (**Fig. 4I**) that can form lateral associations with multiple molecular partners and with each other, organizing the surface membrane proteins in dynamic microdomains ^45^. Tetraspanins promote the entry of multiple viruses, including influenza A virus (IAV), human cytomegalovirus (HCMV), human papillomavirus, hepatitis C virus, Lujo virus, and several alphaviruses ^45^. Depleting the tetraspanin CD9 reduced MERS-CoV lung titers by ∼90% in the infected mice ^47^. Our analyses of ACE2 and TSPAN8 protein expression in the context of SARS-CoV-2 infection of a large panel of lung organoids strongly suggested that TSPAN8 may play a role in SARS-CoV-2 infections. SARS-CoV-2 infection does not lead to TSPAN8 downregulation, whereas other viral infections do. Bioinformatic Analysis using published data performed by the GeneMania ^58^ speculates that ACE2 interacts with TSPAN8 ^59^ and data from Traveglini *et al.* ^3^ showed that epithelial cells are the only cells in the lung that can express both TSPAN8 and ACE2. Alternatively, TSPAN8 may cooperate in the lung organoid infections with other mediators, such as neuropilin-1 (NRP1) described in cell line studies ^56, 57^. Future studies are required to understand the mechanistic details of TSPAN8 and ACE2-mediated SARS-CoV-2 infection.

Clonal cell lines are powerful discovery tools but lack variations in genetic and proteomic traits. Lung epithelial organoids are an attractive platform to study responses to airway infection in the context of diverse cell types ^48^, but every organoid from a human subject is distinct. Our study with a comprehensive panel of lung organoids capitalized on the unique diversity in lung organoids derived from different subjects to discover TSPAN8 as novel biomarker. Furthermore, blocking TSPAN8 showed a reduction of SARS-CoV-2 viral load and TSPAN8 may be a novel potential therapeutic target in SARS-CoV-2 infection.

## Acknowledgements

We thank Drs. Melia Magnen, Leonardo Ferreira, Vilma Arce-Gorvel, Jean-Pierre Gorvel, Jean louis Mege and the entire Roose lab for stimulating discussions. We thank Drs. Hans Clevers, Rob Vries, Bahar Ramezanpour, and Sylvia Boj for helpful discussion on organoid technology. We thank the Aurora Cytek team for technical support on the spectral flow and Arjun Rao for help with scRNAseq analysis. We thank Garcia lab at Standford for the R-spondin surrogates. Most of the work was supported by an Administrative COVID-19 Supplement 3P01AI091580-09S1 (to JPR) on the parent NIH/NIAID P01-AI091580 (Weiss). Organoids were generated in the Roose Organoid D2B unit, started through a UCSF PBBR TMC (Technologies, methodologies, and Cores) grant in 2018 and a gift from the Pathology department and now, in part, funded through a Mark Foundation for Cancer Research Endeavor Program grant (all to JPR). KB is a Mark Foundation Momentum fellow. JRK is funded by the UCSF Bakar ImmunoX Initiative, the UCSF Helen Diller Family Cancer Center, and the American Association for Thoracic Surgery Foundation. DMJ is funded by a private endowment fund. Human Frontier Science Program Fellowship LT000061/2018-L to NKS Additional funds came a grant from the Innovative Genomics Institute and UCSF PBBR (M.O.).

## Disclosure statement

The authors have no potential conflicts of interest.

## Author contributions

LH and JPR: conceived and designed study. LH, SL and KK: performed spectral flow and analyses. LH, KK, SL, OMG: performed organoid experiments. LH and LR: BSL3 work. AS and AJC: scRNAseq experiments and advice. JCL: statistical analyses. LCL: computational analysis of scRNAseq data. LB and DE: advice on antibody staining and lung populations. NKS and MK: H1N1 virus. LH, OMG and KB: microscopy. JZL, VD, SM, MM: patient samples for organoids. GK, DMJ, MM, AJC: (funding) support of their team members. MO: SARS-CoV-2 virus. JRK: surgery lung samples, clinical data discussion. LH, SL, LCL, JPR: figure panels. LH, SL, JPR: manuscript writing. JPR: funding for the study. F, DMJ, MM, MO, MM, AJC, DE, ANS, and JRK: edits on draft manuscript.

---

**Supplemental Figure S1.**
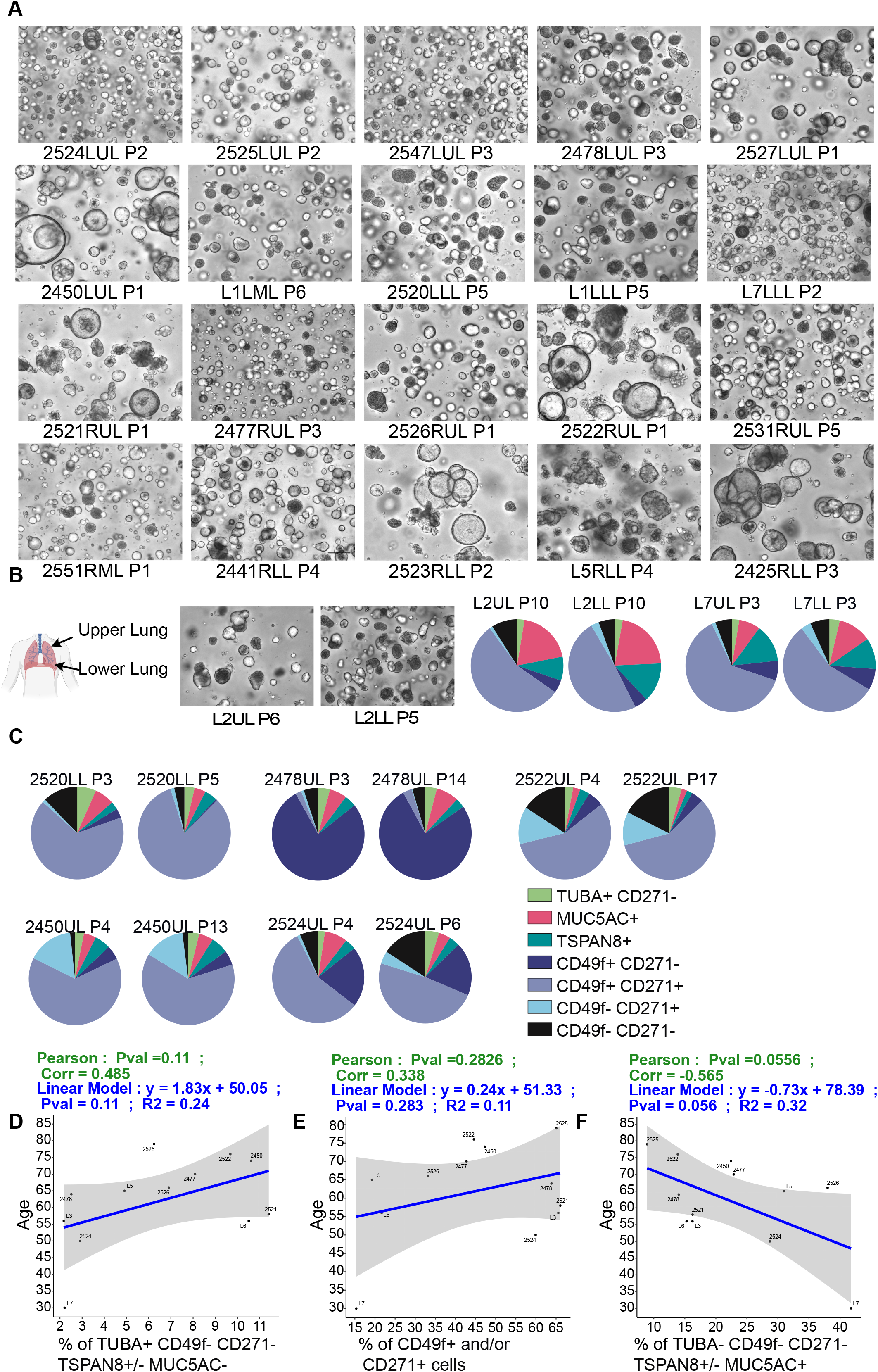

**Supplemental Figure S2.**
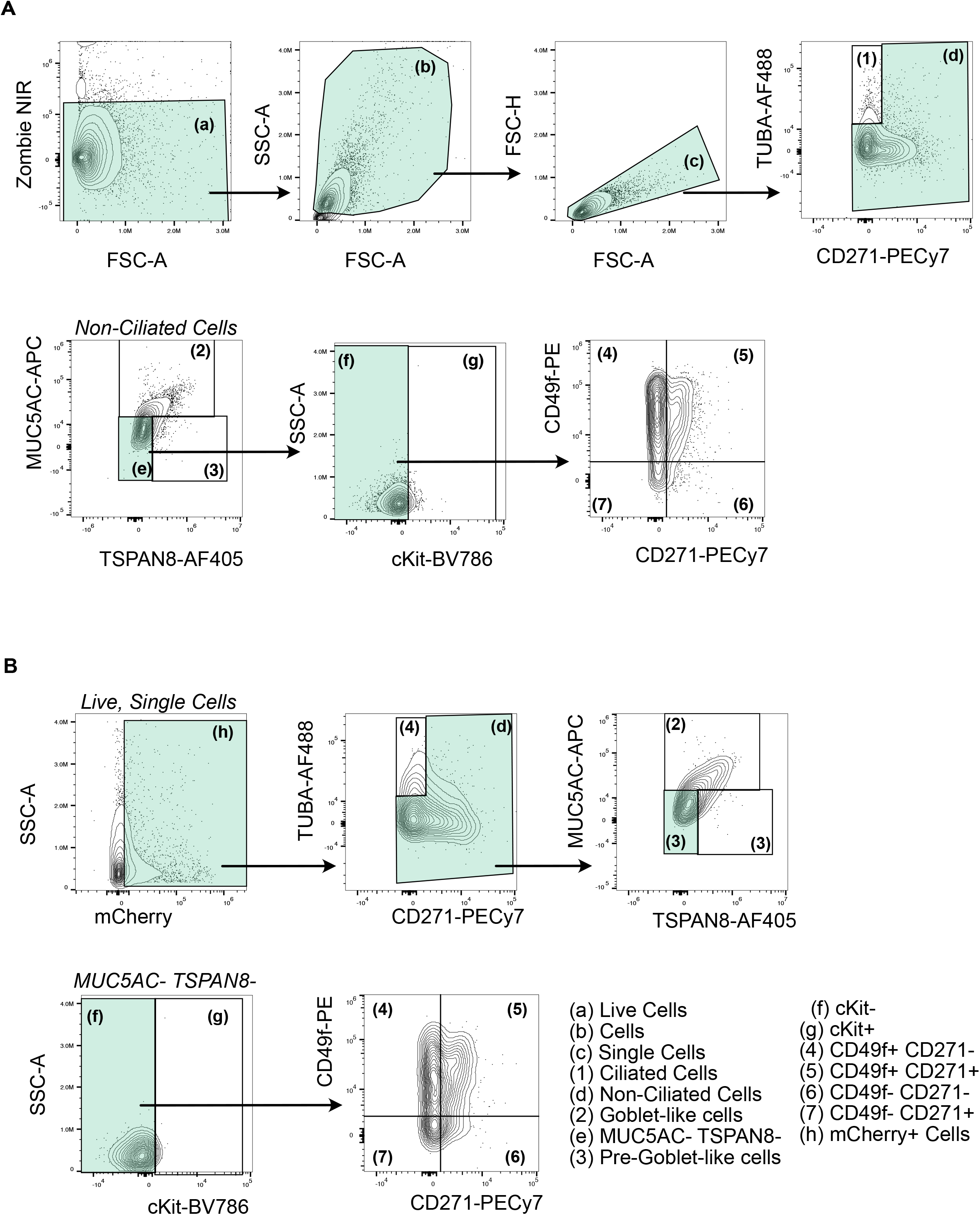

**Supplemental Figure S3.**
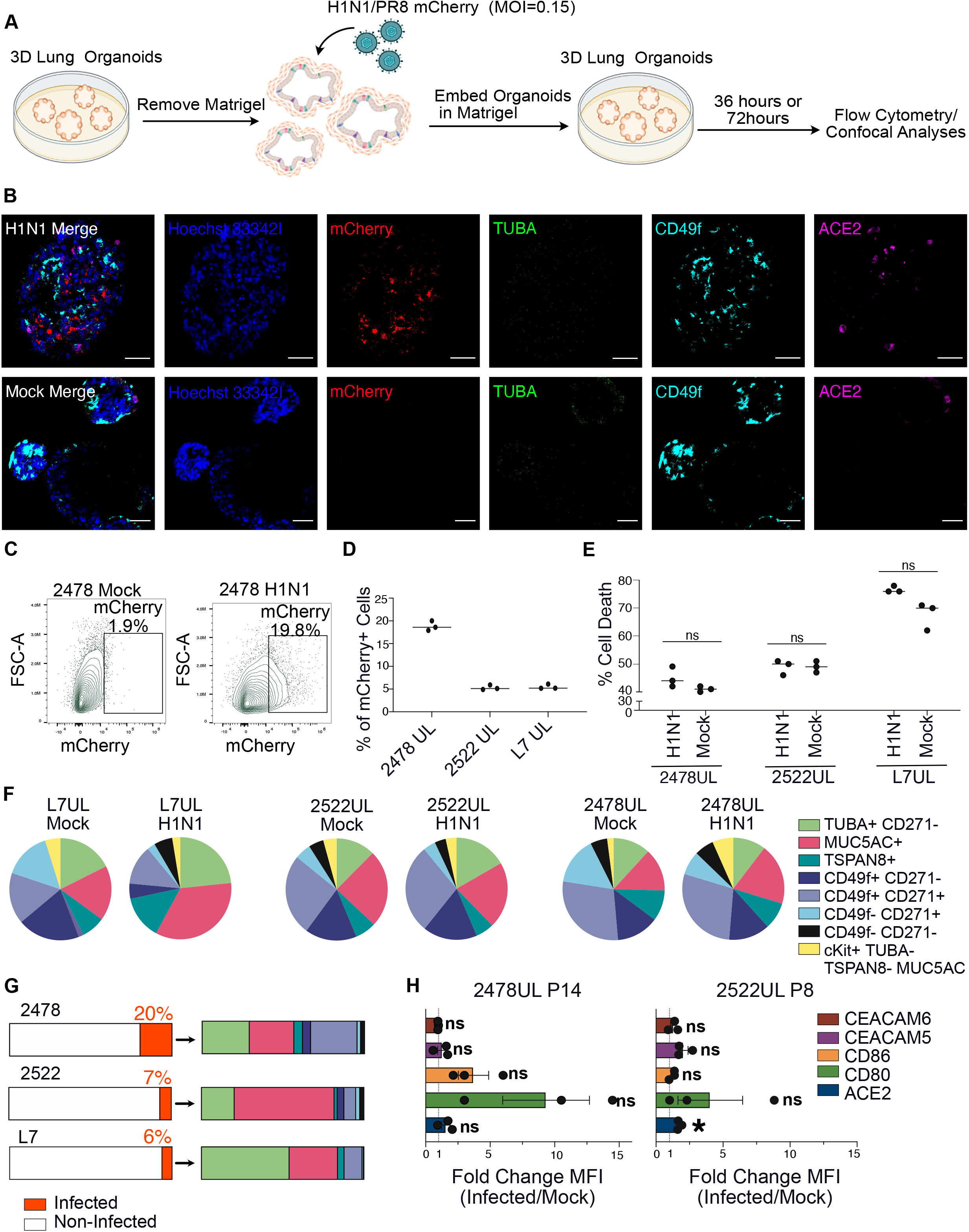

**Supplemental Figure S4.**
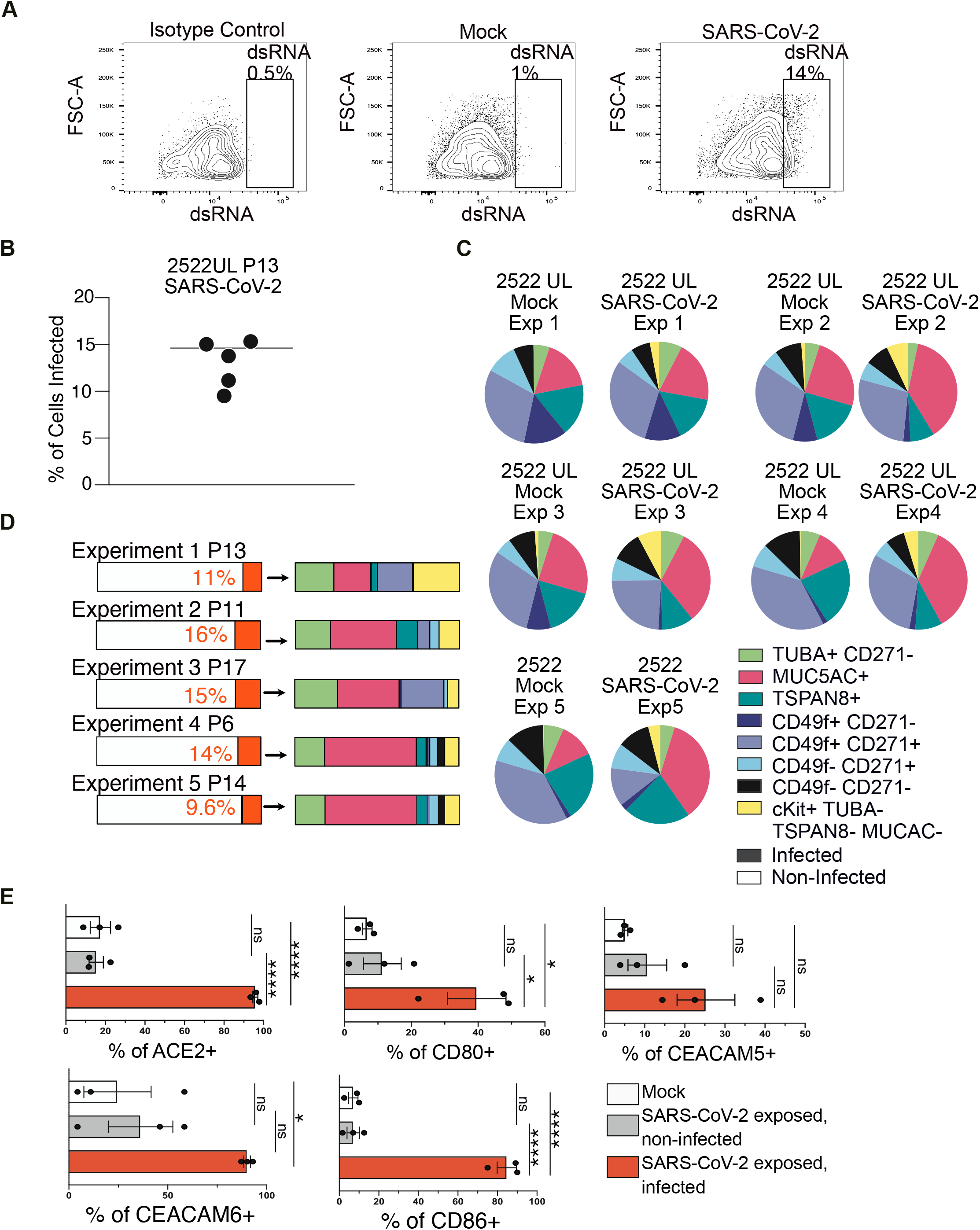

**Supplemental Figure S5.**
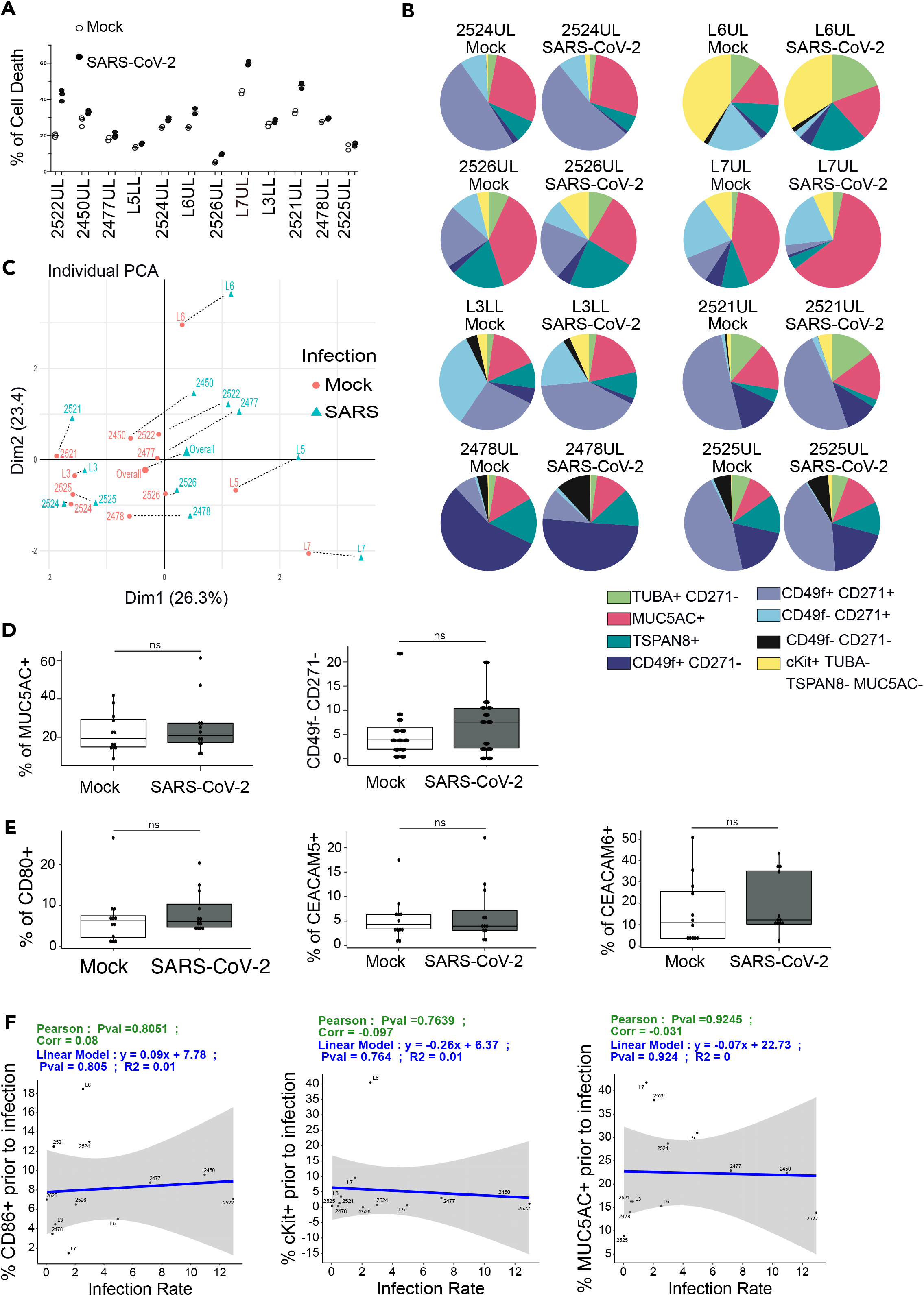

**Supplemental Figure S6.**
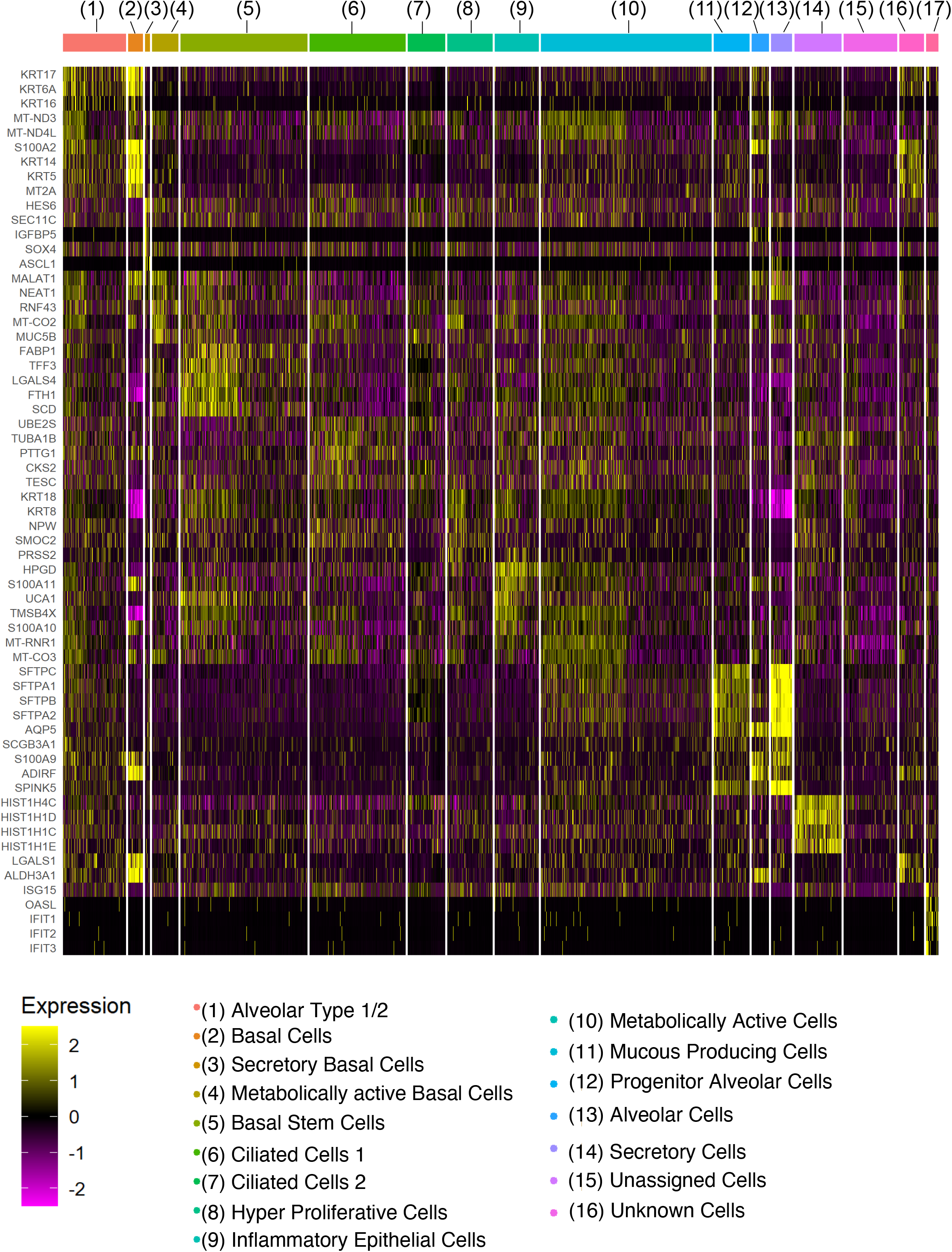

**Supplemental Figure S7.**
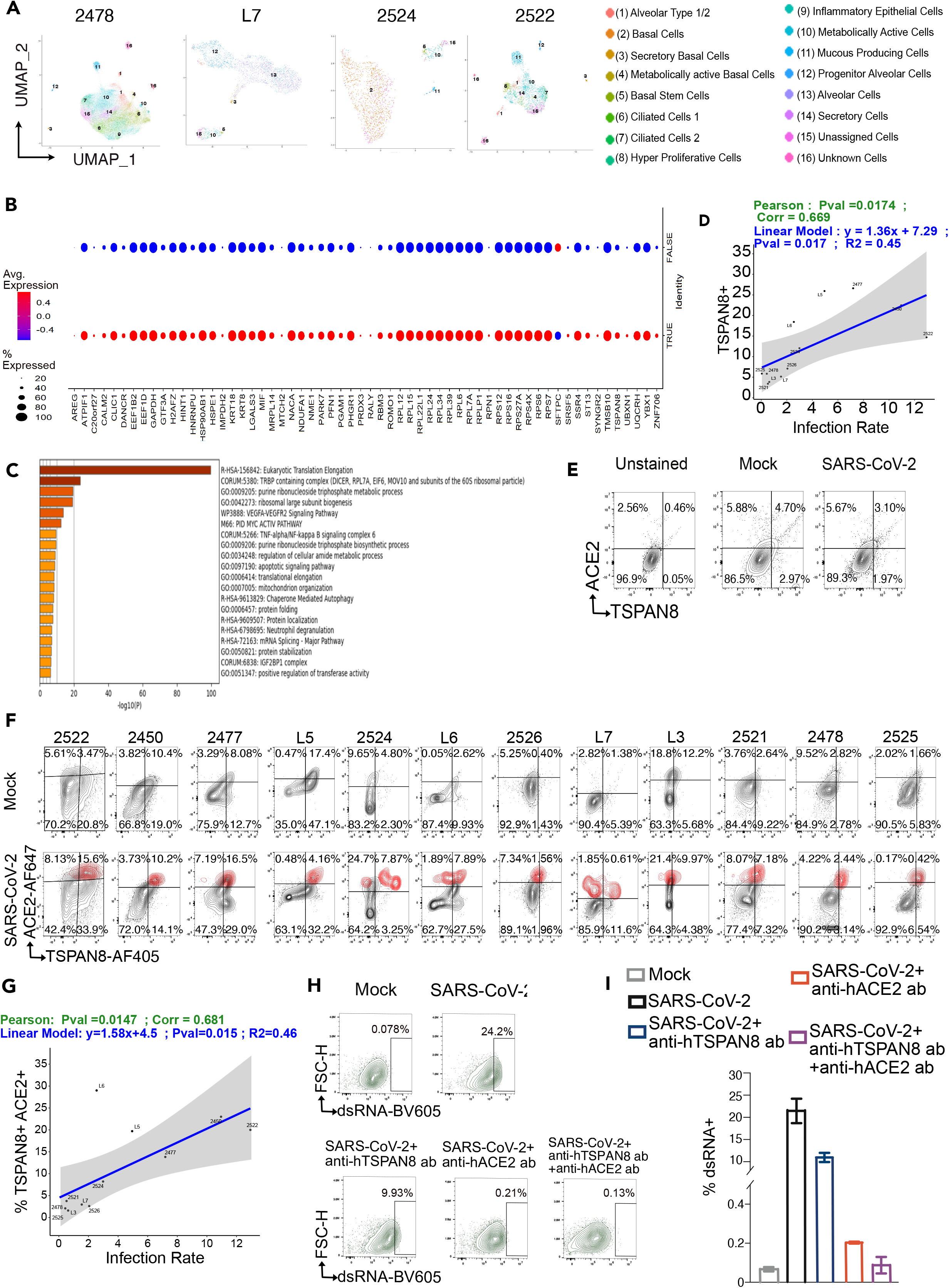

## Notes

### Competing Interest Statement

The authors have declared no competing interest.

