## Supplementary material for "SARS-CoV-2 infection studies in lung organoids identify TSPAN8 as novel mediator": Methods and Supplemental Figure Legends

### Methods and Materials

#### Viruses and Cell Lines

Vero E6 cells were cultured in Dulbecco's Modified Eagle Medium (UCSF Media Production) supplemented with 10% fetal bovine serum (Corning), penicillin/streptomycin (UCSF Media Production) and L-glutamine (Corning) in a humidified incubator at 37C and 5% CO2. SARS-CoV-2 virus (USA-WA1/2020 strain) was propagated in Vero E6 cells. The Vero cells were infected with the SARS-CoV-2 virus, incubated at 37C, 5% CO2 and after 72h the supernatant was collected. The virus was aliquoted and stored at -80C. This work was done under Biosafety Level 3 (BSL-3) conditions.

#### Determination of Virus Titers using Plaque Assay

Viral titer was quantified using a plaque assay in Vero cells. 10-fold dilutions of the virus stock were added to Vero cells in a 12-well plate for 1 hour, after which an overlay of 1.25% Avicel RC-591 in DMEM (UCSF Media Production) was added. The cells were incubated at 37C, 5% CO2 for 72 hours. The cells were fixed with 10% formalin, stained with crystal violet, and washed with water. The plaques were counted to determine the titer of the virus stock. All work was done under BSL-3 conditions.

H1N1/PR8 [1] (gift from Dr. Yoshihiro Kawaoka) was propagated in serum pathogen-free fertilized chicken eggs (Charles River) as previously described [2]. In brief, freshly fertilized eggs were kept in an automatic egg turner for 10 days before injection of the virus into the allantoic cavity. Embryos were incubated with the virus for 2 days. Allantoic fluid was harvested and snap-frozen in liquid nitrogen. Titers were determined with a hemagglutination assay. This work was done under BSL-2 conditions.

#### Patient samples

Human cancer patient samples, 2425LL, 2450UL, 2477UL, 2478UL, 2520LL, 2521UL, 2522UL, 2524UL, 2525UL, 2526UL,2527UL, 2531UL, 2547UL, 2551ML, were collected under UCSF study CC#00654 (IRB 11-06107) from patients undergoing thoracic surgery.

Donor lungs rejected for transplantation were received from organ procurement organization (Donor Network West) as previously described[3]. Explicit approval for the use of donor lungs for research was sought from each donor’s family by Donor Network West as part of the standard organ donation process. Local institutional review board approval is not required because research on tissues from deceased organ donors is not considered human subject research, although institutional biosafety approval was obtained from University of California, San Francisco (UCSF) Institutional Review Board.

**Lung Organoid Culture**

Adult human lung stem cells were derived from non-tumor lung tissue obtained from patients undergoing lung resection or from donor organs. The Roose lab has IRB Exempt certification (IRB #: 12-09467). Isolation of lung stem cells was performed using a protocol adapted from Sachs and colleagues (2019). Lung tissue was washed 2X with DPBS, placed in sterile petri dishes (Corning® Gosselin™) and filled with 10 ml of digestion buffer (DMEM/F12, Collagenase I 1.5mg/ml (Thermo Fischer Scientific, cat#17100017), HEPES (UCSF cell culture facility), penicillin 10,000 IU/mL (Thermo Fischer Scientific Cat#15140122). The tissue chunk was minced in 1-3mm pieces using a scalpel blade and transferred with the digestion buffer into a 50 ml Falcon tube. After 1 hour of incubation at 37Celsius with shaking at 125 rpm, the digested tissue was filtered through a 100µM filter and transferred into a 50 ml Falcon tube. Cells were pelleted (600g, 5 minutes at 4Celsius) and washed twice with DPBS (UCSF Cell Culture Facility). Lung cell pellet was then resuspended in 1300 µL Matrigel (Corning Cat#356230) and plated in ~50 µL droplets in a 24 well tissue culture plate. Plates were placed at 37Celsius with 5% $CO_2$ for 20 min to solidify the Matrigel droplets upon which 550uL of lung organoid media was added to each well. Plates were incubated in the standard tissue culture incubator at 37Celsius. Images were taken on Days 10 to 21 using a BZ-X700 inverted microscope with a CCD cooling camera and BZ-X analysis software (KEYENCE).

**Lung Organoid Biobank**

After 1 to 3 weeks of growth, organoids were dissociated into single cells using TrypLE express. Cells were washed twice with DPBS, pelleted and resuspended in freezing media (Gibco™ Recovery™ Cell Culture Freezing Medium -1560446) and transferred immediately in the Mr. Frosty™ Freezing Container (Cat 51100-001) in the -80C Freezer. Frozen vials were then transferred to the Liquid Nitrogen freezer. When needed, cells were quickly thawed using the 37C water bath, and transferred to 15ml tubes filled with DPBS. Cells were pelleted (600g, 5 minutes at 4Celsius) and washed twice with DPBS. Lung cell pellets were then resuspended in 1300 µL Matrigel (Corning) and plated in ~50 µL droplets in a 24 well tissue culture plate. Plates were placed at 37 degrees Celsius with 5% $CO_2$ for 20 min to solidify the Matrigel droplets upon which 550uL of lung organoid media was added to each well. Plates were incubated in a standard tissue culture incubator at 37 degrees Celsius.

**Lung Organoid Infection**

Lung organoids were removed from Matrigel following 1 minute incubation with Dispase 0.5U/ml (StemCell Technologies, Cat #07913), carefully transferred into a 15 ml tube, washed with DPBS (UCSF Cell Culture Facility) supplemented with 5 mm EDTA (Cellgro, Cat#46-034-CI) and pelleted (200g, 3 minutes at Room Temperature). Organoid pellets were resuspended in 500 µl of lung organoid media. Virus was added at MOI=0.15 for H1N1/PR8 mCherry and MOI=0.3 for SARS-CoV-2. Every organoid well contained 10,000 cells. After 2 hours of incubation, lung organoids were washed twice with PBS and resuspended in 300µL Matrigel (Corning) and plated in ~50 µL droplets in a 24 well tissue culture plate. Plates were placed at 37 degrees Celsius with 5% CO2 for 20 min to solidify the Matrigel droplets upon which 550uL of lung organoid media was added to each well. Plates were incubated in the standard tissue culture incubator at 37 degrees Celsius for 92 hours. After 72 hours spectral flow cytometry analyses (CYTEK Aurora) and Confocal Analyses (Zeiss, SP8) were performed.

**TSPAN8- and ACE2- blocking antibody assays**
Lung organoids were removed from Matrigel following a 1 minute incubation with Dispase 0.5U/ml (StemCell Technologies, Cat #07913), carefully transferred into a 15 ml tube, washed with DPBS (UCSF Cell Culture Facility) supplemented with 5 mm EDTA (Cellgro, Cat#46-034-CI) and pelleted (200g, 3 minutes at Room Temperature). Organoid pellets were resuspended in 500 µl of lung organoid media with human TSPAN8 blocking antibody (R&D Systems, #MAB4734-SP) and/or human ACE2 (R&D Systems, #AF933-SP) at 50µg/ml. After 1 hour incubation with the antibody, SARS-CoV-2 was added. After 2 hours of incubation with the virus, lung organoids were washed twice with PBS and resuspended in 300µL Matrigel (Corning) containing human TSPAN8 blocking antibody (R&D Systems, #MAB4734-SP) and/or human ACE2 (R&D Systems, #AF933-SP) at 50µg/ml. and plated in ~50 µL droplets in a 24 well tissue culture plate. Plates were placed at 37 degrees Celsius with 5% CO2 for 20 min to solidify the Matrigel droplets upon which 550uL of lung organoid media was added to each well. Plates were incubated in the standard tissue culture incubator at 37 degrees Celsius for 24 to 72 hours. Spectral flow cytometry analyses (CYTEK Aurora) and Confocal Analyses (Zeiss, SP8) were performed.

**Flow Cytometry**
Organoids were dissociated into single cells using TrypLE express (ThermoFisher Scientific Cat#12604012). Single Cell suspensions were transferred to 2 ml cryovial tubes and washed twice with FACS buffer (2% FBS, 0,5% BSA, DPBS,0.01% Azide, 0.1mg DNAse + Y-27632 5mM (Abmole) before incubating for 10 minutes with Ig block (True-Stain Monocyte Blocker™ Innovex Bioscience). Cell pellets were incubated with Zoombie NiR (Biolegend) and the antibody mix for 35 minutes followed by two washes with FACS Buffer. Cells were fixed for 30 minutes with Permeabilization Fixation Buffer (Ebioscience). After washing twice with washing buffer cells were incubated for 45

minutes with antibodies staining for intracellular targets followed by a 20-minute incubation with secondary conjugated antibodies. Cells were washed with DPBS before acquiring in the CYTEK Aurora Machine.
Antibodies used for spectral flow cytometry: TSPAN8 AF405, clone 45811(R&D systems), CD66c (CEACAM6) BV510, clone B6.2 (BD Bioscience), EPCAM BV711, clone 9C4, Biolegend; CD86 BV650, clone IT2.2 (BioLegend), CD117 BV785, clone 104D2, BioLegend, CD49f PE, clone GoH3 (BioLegend), CD80 PE-Cy5, clone 2D10 (BioLegend), CD271 PE-Cy7, clone ME20.4, BioLegend, ACE2 AF647, clone Q9BYF1 (R&D systems), CEACAM5 APC-Fire750, clone 487609 (R&D systems), MUC5AC Biotin, clone 45M1 (ThermoFischer Scientific), Streptavidine APC (eBioscience), anti-TMPRSS2, clone EPR3862 (Abcam), acetylated Tubulin AF488, clone 6-11B-1 (SantaCruz Biotechnologies), anti-dsRNA (Scicons), IGg BV605, clone4053 (Biolegend)

**Confocal Imaging**

Lung organoids were removed from Matrigel following 1 minute incubation with Dispase 0.5U/ml (StemCell Technologies, Cat #07913), carefully transferred into a 15 ml tube, washed with DPBS (UCSF Facility) and pelleted (200g, 2 minutes at Room Temperature). The organoid pellet was incubated for 30 minutes in 4% PFA 10% FBS Triton 0.1X and TrueStain. Organoids were washed twice with PBS-Triton 0.1X and stained overnight at 4C with primary antibodies dsRNA Ab (clone J2, Scicons), SARS-CoV-2 Nucleocapsid Ab (Novus Biologicals, Cat#NB100-56576SS). Organoids were then washed and incubated for two hours with secondary antibodies Goat anti mouse IGg AF555, clone4053 (Biolegend), CD49f PE, clone GoH3 (BioLegend), ACE2 AF647, clone Q9BYF1 (R&D systems), and acetylated Tubulin AF488, clone 6-11B-1 (SantaCruz Biotechnologies). After staining, organoids were washed with DPBS- 0.1X Triton and resuspended in Fructose-Glycerol Clearing Solution 60% (vol/vol) glycerol and 2.5 M fructose. Organoids were mounted on coverslips and imaged with Leica SP8 Confocal Microscopy. Confocal Z-stack images were generated using the staining maxima.

**Statistics**

Statistical analyses were run in R (version 4.0.2) (quote 1). Paired Samples Wilcoxon Test were performed using Wilcox.Test (stat v4.0.2) (quote 1). Outliers were removed if the value was over Q3+1.5 IQR. We used PCA (FactoMineR v2.4) to perform Principal Component Analysis and fviz_pca_ind or fviz_pca_var (factoextra v1.0.7) (quote 3) for visualization. Community distances were evaluated based on relative abundance of cell populations by PERMANOVA (Bray, *p<0.05). For spearman's and pearson's correlations, we used cor function (stat v4.0.2) to compute the correlation coefficients and lm (stat v4.0.2) to fit linear models. Data management was done using tidyverse (v1.3.0) (quote 4). All graphs were built using ggplot2 (quote 5) [4, 5, 6, 7, 8].

UMAP dimensionality reduction, discovery of upregulated/downregulated genes, and gene expression related plots were constructed in R v 4.0.3 (1) via Seurat v 4.0 provided by the Satija Lab [9] and ggplot2 v 3.3.3 Clusters were separated using the Louvain clustering method with a resolution of 0.6, and upregulated differential expression gene scores between clusters were used to establish cell type identities[8].
Bar graph of enrichment analysis up regulated pathway and processes based in cells positive for SARS-CoV-2 reads were generated using Metascape[10].

**Software**
GraphPad Prism 9 was used for DATA visualization. Fiji/ImageJ (V) software 2.1.0/1.53c was used for confocal microscopy analysis. FlowJO and CYTEK Aurora software 10.7.1 was used for spectral flow analyses.

**Single cell and library preparation for scRNA-sequencing**

For the single-cell RNA sequencing experiments each organoid was generated from 4 different donors. After organoid dissociation into single-cell suspension using TrypLE express (ThermoFisher Scientific Cat#12604012), cells were counted, and pooled together (78,000 total cells). Pooled cells were then loaded evenly across two lanes in the Chromium Controller for generating single-cell contained in lipid droplets following the manufacturer's instructions (10X Genomics). A Chromium Single cell 3' Reagent Kit (v3.1) (10X Genomics) was used for reverse transcription, cDNA amplification and library construction of the gene expression libraries following the manufacturer's instructions. All samples were encapsulated, and cDNA was generated within 6 hours after organoid processing. Finally, Pooled libraries were sequenced on an Illumina NovaSeq 6000.

**Single cell RNAseq analysis**
**Data pre-processing of 10x Genomics Chromium scRNA-seq data:**
Data pre-processing was performed as previously described (CITE: Combes et al, Nature, 2021). Briefly, sequencer-obtained bcl files were demultiplexed into individual samples using the Cellranger 3.0.2 suite of tools (https://support.10xgenomics.com). Feature-barcode matrices were obtained for each sample by aligning the raw fastqs to GRCh38 reference genome (annotated with Ensembl v85) using the Cellranger count. Raw feature-barcode matrices were loaded into Seurat and genes with fewer than 3 UMIs were dropped from the analyses. Matrices were further filtered to remove events with greater than 30% percent mitochondrial content, events with greater than 50% ribosomal

content, or events with fewer than 250total genes. The cell cycle state of each cell was assessed using a published set of genes associated with various stages of human mitosis.

**Inter-sample doublet detection:**
Inter-sample doublet detection was performed as previously described (CITE: Combes). Libraries containing samples pooled prior to loading were processed using Freemuxlet (https://github.com/statgen/popscle) to identify clusters of cells belonging to the same patient via SNP concordance. Cells are classified as singlets arising from a single library, doublets arising from two or more libraries, or as ambiguous cells that cannot be accurately assigned to any existing cluster (due to a lack of sufficient genetic information).

**Data quality control and Normalization:**
The filtered count matrices were normalized, and variance stabilized using negative binomial regression via the scTransform method offered by Seurat [11]. The effects of mitochondrial content, ribosomal content, and cell cycle state were regressed out of the normalized data to prevent any confounding signal. The scTransformed data from different sequencing libraries were combined and normalized using Harmony sc integration software.

**Intra-sample heterotypic doublet detection:**
All libraries were further processed to identify heterotypic doublets arising from the 10X sample loading. Processed, annotated Seurat objects were processed using the DoubletFinder package [12]. Briefly, the cells from the object are modified to generate artificial duplicates, and true doublets in the dataset are identified based on similarity to the artificial doublets in the modified gene space. The prior doublet rate per library was approximated using the information provided in the 10x knowledgebase (https://kb.10xgenomics.com/hc/en-us/articles/360001378811) and this was corrected to account for homotypic doublets using the per-cluster numbers in each dataset.

**Differential expression tests and cluster marker genes, cluster annotation:**
Differential gene expression (DGE) tests were performed on log-normalized gene counts using the Poisson test (with a latent batch variable to account for multiple library preparations) provided by the FindMarkers/FindAllMarkers functions in Seurat. Cluster marker genes were identified by applying the DE tests for upregulated genes between cells in one cluster to all other clusters in the dataset. Top ranked genes (by log-fold changes) from each cluster and between clusters are used for further analysis.

**Data Mining**

To examine presence of *ACE2* and *TSPAN8* co-expressing cells in vivo in normal human lung, we analyzed the processed 10X single-cell sequencing data from Travaglini et al[13]. Of the 60,993 cells derived from lung tissue of 3 patient donors in this dataset, 48 cells were found with at least 1 UMI (unique molecular identifier) for both genes. Only 1 cell derived from patient 1, which had fewer cells sequenced overall and so we excluded it. The expression values represent ln(UMI-per-10K + 1) in each of the 47 cells from patients 2 and 3. Cell type designations were determined by Travaglini *et al*.

Differential expression of TSPAN8 in nasal swabs of adult patients with acute respiratory illness (ARI) due to COVID-19 (n=93) or other viral infection (n=41), in comparison to patients with ARI due to non-viral etiology (n=100), was derived from Mick et al[14]. The differential expression analysis between the 3 viral status groups was performed with the R package limma while controlling for gender and age.

**Legends Supplementary Figures**

**Supplementary 1: Donor derived lung organoids display distinct phenotypes.**

A) Brightfield images of twenty lung organoids derived from different donors.
B) Brightfield images of organoids derived from the upper and lower region of the left lung of individual L2. (Right) The composition of lung organoids based on spectral flow cytometry analyses derived from the upper and lower lobes of subject L2 and L7 are represented in pie charts. Each part of the pie chart represents the mean value of the percentage of the cells population in distinct donor derived lung organoids.
C) Lung organoids compositions derived from spectral flow cytometry analysis of different passages of organoids from multiple donors. Each section of the pie chart represents the cell population mean from 3 independent experiments.
(D-F) Linear regression showing the relationship between age of the donor and cell types found in the donor derived lung organoids (N=3, n=3). Each dot represents the mean value of the percentage of the cells population in distinct donor derived lung organoids. Values for Pearson correlations and corresponding P values are depicted. Functions of the positive or negative correlations are depicted by the Linear Model.
(D) Age and % of ciliated like cells (TUBA+ CD271-). (E) Age and % of basal stem cells (CD271+ or CD49f+). (F) Correlation between age of the donors and proportion of mucus producing cells.

**Supplementary Figure 2: Spectral flow cytometry analyses of mock and infected lung organoids.**

A) Spectral flow cytometry gating strategy. Cell populations are defined after excluding ZombieNIR-positive, dead cells and doublets in steps a-c. (1) TUBA + CD271- are considered as ciliated cells. Fraction (d) is analyzed for MUC5AS and TSPAN8 and (2) MUC5AC+ TUBA- cells are considered goblet-like cells. (3) TSPAN8+ MUC5AC- TUBA- are defined as pre-goblet cells. Fraction (e) is analyzed for cKit and (g) are considered cKit-positive cells. Fraction (f) is analyzed for CD49f and CD271, (4) CD49f+ CD271-, (5) CD49f+ CD271+ and (6) CD49f- CD271+ are considered basal stem cells and (7) CD49f- CD271- are undefined cells.

B) Spectral flow cytometry gating strategy for mCherry+ cell populations. The same gates as in Supplementary 1 A) were drawn on live, single cell and mCherry+ cells (h).

**Supplementary Figure 3: H1N1 viral infection of lung organoids.**

A) Experimental scheme of lung organoid infection with H1N1/PR8 mCherry virus.

B) Confocal images (Z-stack) of H1N1 infected (top panel) and mock infected (bottom panel) whole mounted lung organoids. Scale bars are 50uM.

C) Spectral flow cytometry plot of live cells and gating for mCherry-positive cells.

D) Percentage of H1N1/PR8 mCherry-positive lung organoids.

E) Percentage of cell death in indicated organoids.

F) Cell composition in mock and infected organoids. Pie charts represent mean percentages of n=3 independent experiments for each donor derived organoids.

G) Percentage of H1N1/PR9 mCherry+ cells (left) and H1N1/PR8 mCherry-infected cell types (right). Colors as in 2F. Mean values of n=3 experiments shown in bars.

H) Fold change in median fluorescence intensity (MFI) of CEACAM6, CEACAM5, CD80, CD86 and ACE2 during infection at 36h H1N1 p.i. (MOI=0.15). Bars represent mean, error bars are SEM, n=3. Paired t-test, *$p < 0.05$; ns, non-significant.

**Supplementary Figure 4. Reproducibility of SARS-CoV-2 infection in lung organoids.**

A) dsRNA staining for spectral flow cytometry analyses of SARS-CoV-2 infected organoids at 72h p.i.
B) Percentage of dsRNA+ in replicate experiments infecting the same donor derived organoid with SARS-CoV-2. Each point represents an independent experiment.
C) Cell composition of organoid 2522UL in Mock and infected organoids at 72h p.i. (MOI=0.3) in five separate experiments.
D) The percentage of dsRNA+ cells (left) and the cell types in dsRNA+ cells (right) based on spectral flow cytometry analyses.
E) The percentage of cells positive for CEACAM6, CEACAM5, CD80, CD86 and ACE2 at 72h p.i. in mock infected, SARS-CoV-2 exposed (dsRNA-) and SARS-CoV-2 infected (dsRNA+) cells from organoid 2522UL. Every point represents data for a single experiment. Bars represent mean, error bars are SEM. Paired t-test, $*p < 0.05$, $****p<0.0001$; ns, non-significant.

**Supplementary Figure 5: Donor derived lung organoid response to SARS-CoV-2.**

A) The percent of cell death in mock and SARS-CoV-2 infected organoids (MOI=0.3, 72h p.i.). Each DOT represents the mean value of the percentage of the cells population in distinct donor derived lung organoids (N=3, n=3).
B) Cell composition based on spectral flow cytometry analyses of mock and SARS-CoV-2 infected organoids (MOI=0.3, 72h p.i.). Each part of the pie chart represents the mean value of the percentage of the cells population in distinct donor derived lung organoids (N=3, n=3).
C) Principal component analysis using cell populations and infection conditions. PCA reduced the descriptors into two dimensions as shown in the individual plot. Each point corresponds to an observation in the dataset and were colored according to their mock or infection state. Community distances were evaluated based on relative abundance of cell populations by PERMANOVA (Bray, $*p<0.05$).
D) Comprehensive analyses of the percentage of MUC5AC+ and CD49f- CD271- cells in mock and infected organoids. Each point represents the mean of % cell type for the distinct donor for Mock or SARS-CoV-2 conditions. Wilcoxon signed-rank test, $*p<0.05$, N=12, n=3. Outliers were removed if the value was over Q3+1.5 IQR.
E) Comprehensive analyses of the percentage of cells positive for activation markers in mock and infected organoids. Wilcoxon signed-rank test, $*p<0.05$, N=12, n=3. Outliers were removed if the value was over Q3+1.5 IQR.
F) Linear regression showing the relationship between SARS-CoV-2 infection rate and fraction CD86+ cells, TUBA+ cells and MUC5AC+ cells prior to infection. Each dot represents the mean of % cell type in 3 independent experiment (n=3) for the distinct donor. Linear regression, R2 value and statistical significance stated on graph. Mean values, N=2 to 3, n=3 for every donor derived organoid.

**Supplementary Figure 6: Differentially expressed genes donor derived organoids.**

Clustermap of top differentially regulated genes between clusters established in Fig 5A. Hierarchical clustering is used on a concatenated set of genes constructed from the top 5 statistically significant expressed genes from each cluster.

**Supplementary Figure 7: TSPAN8 a potential biomarker in COVID-19 severity.**

A) UMAP's constructed for individual organoids from four different subjects. cell subsets from the merged data, with overlaid clusterings from Fig 5A.
B) Top statistically significant genes when computing differential expression between cells that had any genes mapping to the covid genome (TRUE) versus those that had no genes mapped (FALSE). The size of the dot shows the percentage of cells expressing the gene.
C) Bar graph of enrichment analysis up regulated pathway and processes based on cells positive for SARS-CoV-2 reads, colored by p-values. The darker the color, the lower the p-value. Figure created in Metascape.
D) Analysis of cell populations in mock and infected organoids. Percentages of TSPAN8+ cells. Wilcoxon signed-rank test, $*p<0.05$, N=12, n=3.
E) Example of gating strategy of ACE2 and TSPAN8 expression. For all the organoids and the individual staining sets, quadrants were defined on the basis on unstained controls.
F) Spectral flow cytometry plots showing live dsRNAneg cells (in gray) and live dsRNApos cells (in red- overlaid) in Mock and SARS-CoV-2 infected organoids at 72 hours p.i. (MOI=0.3).
G) Linear regression showing the relationship between the fraction of TSPAN8+ ACE2+ cell population in SARS-CoV-2 infected organoids and infection rate. Linear regression, R2 value and statistical significance stated on graph. Mean values, N=12, n=3.

H-I)Flow Cytometry plots (H) and graph of Spectral Flow Cytometry analysis of dsRNA+ cells (I) in 2450UL lung organoids, 24 hours post SARS-CoV-2 infection (MOI=0.3), in absence or presence of TSPAN8 or ACE2 blocking antibodies (n=3). Bars represent mean, error bars are SEM. Paired t-test, $*p < 0.05$, $****p<0.0001$; ns, non-significant.

**Supplementary Table 1:**
Organoid Donor Records
HTN= hypertension, HLD= hypersensitive lung disease, Afib= atrial fibrillation, CAD= coronary artery disease.

**Supplementary Table 2:**
List of genes expressed in different clusters of the UMAP in Figure 5A

**Supplementary Table 3:**
Differentially expressed genes coding for cell receptors in cells negative and positive for SARS-CoV-2 genome reads.
