## Supplementary material for "SARS-CoV-2 infection studies in lung organoids identify TSPAN8 as novel mediator": Table 1

| Sample ID | Age | Sex | Smoker (Y/N) | Heart Condition (Y/N) | Other Conditions | Location of Sample | Lung Sample Side | Specific Sample Location |
|---|---|---|---|---|---|---|---|---|
| 2425LL | 81 | F | Y | Y | NSCLC, CAD, HTN, HLD | Lower Lobe | Right | |
| 2441LL | 49 | F | N | N | NSCLC | Lower Lobe | Right | |
| 2450UL | 74 | F | Y | Y | NSCLC, HTN, HLD, Non-Sustained and Paroxysmal Ventricular Tachycardia (on beta-blocker therapy) | Upper Lobe | Left | |
| 2477UL | 70 | F | Y | N | NSCLC | Upper Lobe | Right | |
| 2478UL | 64 | F | Y | Y | NSCLC, HTN, HLD, Afib (on Xarelto) | Upper Lobe | Left | |
| 2520LL | 46 | F | N | N | NSCLC | Lower Lobe | Left | Anterior Basilar Segment |
| 2521UL | 58 | M | Y | Y | NSCLC, Afib | Upper Lobe | Right | |
| 2522UL | 76 | F | Y | N | NSCLC, HTN | Upper Lobe | Right | |
| 2523LL | 38 | F | N | N | NSCLC, Unconfirmed Tachycardia | Lower Lobe | Right | |
| 2524UL | 50 | F | N | N | NSCLC | Upper Lobe | Left | |
| 2525UL | 79 | F | N | N | NSCLC | Upper Lobe | Left | Apical |
| 2526UL | 66 | F | Y | N | NSCLC | Upper Lobe | Right | |
| 2527UL | 70 | F | Y | N | NSCLC | Upper Lobe | Left | |
| 2531UL | 80 | M | Y (former) | N | NSCLC | Upper Lobe | Right | |
| 2547UL | 76 | F | Y | N | NSCLC, CAD, HTN | Upper Lobe | Left | |
| 2551ML | 26 | F | Y | N | NSCLC | Middle Lobe | Right | |
| L1UL, L1ML, L1LL | 32 | M | Y (meth, marijuana) | N/A | Psychaitric Problems | Upper, Middle, Lower Lobe | Left | |
| L2UL, L2LL | 27 | M | Y | N/A | N/A | Upper, Lower Lobe | Left | |
| L3LL | 56 | M | N | N/A | Bronchial Asthma | Lower Lobe | Left | |
| L5LL | 65 | F | Y | N/A | Post-Op Pnemonia on Emphysema/ Bronchoectasia Background | Lower Lobe | Right | |
| L6UL | 56 | F | N | N/A | N/A | Upper Lobe | Left | |
| L7UL, L7LL | 30 | M | Y (former) | N/A | Asthma, Diabetes, HTN | Upper, Lower Lobe | Left | |
